# Complex landscapes partially mitigate negative pesticide effects on tropical pollinator communities

**DOI:** 10.1101/2021.03.10.434664

**Authors:** Diana Obregon, Olger R. Guerrero, Elena Stashenko, Katja Poveda

## Abstract

Land-use change and pesticides have been identified as two of the main causes behind pollinator decline. Understanding how these factors affect crop pollinator communities is crucial to inform practices that generate optimal pollination and ensure sustainable food production. In this study, we investigated the effects of landscape composition and pesticide residues on bee communities and their pollination services in *Solanum quitoense* “lulo” crops in Colombia. On 10 farms, located along a gradient of landscape complexity that varied from 0.15 to 0.62 in their natural habitat proportion, we characterized the bee community visiting the crop, and carried out pollination experiments with bagged and open inflorescences to later estimate fruit set, weight, and diameter at every site. Additionally, we performed pesticide analysis on collected anthers through liquid chromatography to estimate pesticide risk coming from the crop fields using hazard quotients (HQ). Bee abundance and species richness decreased with increased HQ, but these negative pesticide effects were less detrimental in farms with higher natural habitat proportions. However, this buffer effect was lost at sites with very high HQs. Imidacloprid was frequently found in the anthers and there were extremely high concentrations in some farms (0.6 to 13063 μg/kg), representing the molecule of greater risk for bees in this context. Pollinator’s importance to crop yield was demonstrated in the exclusion experiments, where we found a reduction in fruit set (51%), weight (39%), and diameter (25%). We found a significant effect of bee richness on fruit set, while landscape composition and HQ had no significant effect on fruit set, suggesting that the last two factors do not affect yield directly, but indirectly through a decrease in pollinator diversity. Our results provide novel evidence that the natural habitat loss due to the expansion of pastures for cattle ranching and pesticide residues in anthers reduce bee diversity and abundance in this Andean cropping system, but strategies to protect and restore natural habitat can help to buffer, until certain levels, these negative effects.

**Highlights:**

- We explored how landscape composition and pesticide residues impact bee communities and pollination services in *Solanum quitoense* crops.
- As the proportion of natural habitat in the landscape increased, bee richness also increased. While as pesticide hazard quotients in *S. quitoense* anthers increased, bee diversity and abundance decreased.
- The natural habitat surrounding farms mitigates the negative pesticide effects on bees when hazard quotients are low/medium, but not when they are high.
- *S. quitoense* crops are highly dependent on bees for optimum yield.

## 1. Introduction

Crop pollination is a fundamental ecosystem service that provides significant benefits to about 75% of the leading global crops (Klein et al., 2007). Vegetables, fruits, nuts, and edible oil crops are among the most pollinator-dependent crops (Gallai et al., 2009). Insect pollination can increase fruit and seed set but also quality variables such as weight, symmetry, and nutritional value (Bartomeus et al., 2014; Bommarco et al., 2012; Garratt et al., 2014). Although there is a great number of insect taxa contributing to crop pollination (Rader et al., 2016), bee abundance and diversity are key to achieve optimum yields (Garibaldi et al., 2016; Winfree et al., 2018).

There is evidence that bee populations are declining (Bartomeus et al., 2013; Cameron et al., 2011; Mathiasson and Rehan, 2019). Land-use change, limited foraging resources, bee pathogens, pesticides, and the interaction among these factors have been identified as the main potential causes (Goulson et al., 2015). At the landscape scale, increases in cultivated areas are associated with natural habitat reduction and fragmentation (Petit and Firbank, 2006), leading to less floral and nesting resources for bees (Heard et al., 2007; Potts et al., 2005) and to lower bee abundance and diversity, with negative impacts on crop pollination services (Bommarco et al., 2013; Connelly et al., 2015; Kremen et al., 2002; Le Féon et al., 2010).

In agricultural settings, bees are frequently exposed to pesticides through direct contact while foraging or digging into the soil, or through the consumption of nectar, pollen, and water with pesticide residues (Krupke et al., 2012; Stoner and Eitzer, 2012). For example, neonicotinoids, are systemic insecticides commonly used in agriculture as foliar sprays, soil drench, or seed dressings to control a wide range of pests (Goulson, 2013). These insecticides, such as imidacloprid, are highly toxic for bees, and represent an important threat to pollinator communities due to their frequent application (Sanchez-Bayo and Goka, 2014). In laboratory conditions, it is widely demonstrated that neonicotinoids have lethal and sublethal effects on bees, such as associative learning reduction, a decrease in foraging activity (Blacquière et al., 2012) and a reduced immunocompetence (Aufauvre et al., 2012; Brandt et al., 2016). While in the field, some studies have shown negative effects of neonicotinoids on bees such as reduced colony growth in bumble bees (Goulson, 2015) or a reduced wild bee density and solitary bee nesting (Rundlöf et al., 2015), others have not found any effects (Pilling et al., 2013, Cutler et al., 2014). These inconclusive results indicate that more field experiments are needed to understand how realistic pesticide exposure affects pollinator communities in different contexts (Lundin et al., 2015).

Although the effects of landscape simplification and pesticides on bees are relatively well established, the interaction of these two stressors has not been significantly explored. In apple orchards in New York, it was shown that the use of pesticides is negatively correlated with bee richness, but this effect was weakened on farms with more natural habitat in the landscape (Park et al., 2015). In India, a study looking at bee diversity found that both semi-natural habitat areas, as well as pesticide use, regulate species diversity at the landscape scale (Basu et al., 2016). However, those studies are based on pesticide use reported by farmers and actual pesticide residues in the field were unknown.

Most of the work investigating the drivers that cause pollinator decline has been conducted in Europe and the US and it is mostly focused on honeybees and bumblebees (Lundin et al., 2015). For tropical regions, where the agricultural landscape is very heterogeneous and comprised of small landholdings and greater crop diversity, the information is limited (Brosi et al., 2007, 2008), making it difficult to develop strategies for conservation and restoration (Winfree et al., 2008). A case in point are the Andean mountains, which are experiencing a rapid loss of natural habitats due to their transformation into cropland and grazing areas (Etter et al., 2006; Foley et al., 2005). Pastures are now the main landcover type in the Andes, representing 65.2% of the total area and the biggest driver of forest losses (Rodríguez Eraso et al., 2013). The tendency to increase pesticide use in developing countries without the regulation of harmful molecules and consideration for effects on biodiversity is troubling (Lundin et al., 2015; Schreinemachers and Tipraqsa, 2012). So far, in this region, there is little information that shows the effects of this accelerated land transformation or pesticide use on the local wild bee populations or their effects on pollination services. Therefore, this study aimed to establish how both landscape composition and pesticide residues affect the bee community and the pollination services in a traditional Andean crop (*Solanum quitoense* “Lulo”). Specifically, we were interested in testing if natural areas can actually buffer against the negative effects of pesticide exposure on bees in these Andean systems. To accomplish this goal, we characterized the flower visitors and the crop pollinator dependency along a landscape simplification gradient. We also screened for pesticide residues in lulo flower anthers to estimate the risk that this crop represents for the pollinator community.

## 2. Methods

### 2.1 Study system

*Solanum quitoense* (Solanaceae), known as “lulo” or “naranjilla”, is an Andean fruit used in Colombia and Ecuador for fresh consumption and by the juice industry (Sánchez Fory et al., 2010). Lulo plants are self-compatible, and strongly andromonoecious, producing both hermaphroditic and staminate flowers in the same inflorescences. Hermaphrodite flowers are distinctive by their long styles that exceed the anthers (Miller and Diggle, 2003). *Solanum* flowers produce no nectar and rely on the production of abundant protein-rich pollen to reward pollinators (Buchmann, 1986). Anthers are poricidal and require sonication to release the pollen (Buchmann and Cane, 1989). Palynological identification of pollen provisions have shown that lulo pollen is collected by *Bombus pullatus, Bombus atratus* (Mercado et al., 2017), and *Euglossa nigropilosa* (Otero et al., 2014). Greenhouse experiments demonstrated the efficient use of commercial colonies of *Bombus atratus* (Almanza, 2007), and *Bombus terrestris* (Messinger et al., 2016) as pollinators but there is no information on the wild pollinator community.

### 2.2 Study region and landscape composition

This study was conducted in 2018 in the municipality of Chameza (Casanare) on the eastern slope of the eastern Andean Cordillera in Colombia. The study area is dominated by forested mountains with steep slopes (Quintana et al., 2017), with all our sites located in elevations between 1188 and 1683 m. Current land covers in the area are primary and secondary forest (called natural habitat in this paper), pastures (grazing lands composed mostly of species of the families Poaceae and Cyperaceae), and small-scale agricultural areas (mostly lulo and sugarcane). In July of 2018, we selected 10 farms with conventional lulo production. Farms were separated by a minimum distance of 1.4 km up to 15 km, and the areas planted with lulo varied from 1200 to 7100m^2^. Using a DJI Phantom 4 drone, and the Map Pilot application, we acquired high-resolution images (5cm/pixel) at 500 m around the center of every lulo crop. The images per site were combined in orthophotos in which polygons of the different land cover types (natural habitat, pastures, and agriculture at the 500 m radius) were manually drawn to calculate the area, and the different land covers’ proportion using QGIS 2.18 Las Palmas (2016). Among the land cover types, the proportion of natural habitat and the proportion pasture were highly correlated *(Pearson’s r = −0.97, p<0.001, n=10),* and together made up the majority of the area on each farm (87% to 99%). Based on this close correlation, we used natural habitat as the landscape explanatory variable. The natural habitat proportion among farms ranged from 0.15 to 0.62, forming a landscape gradient from simple to complex landscapes respectively (Supplementary Figure S1 and Table S2).

### 2.3 Pesticide Analysis

We collected all the anthers of one flower from 10 different plants at every farm to assess the pesticide exposure risk of bees in the crop. The pesticide analysis was performed in the Laboratory of chromatography and mass spectrometry CROM-MASS at the Industrial University of Santander. Samples were tested for the nine most commonly used pesticides in the crop (Insecticides: Methomyl, Abamectin, Bifenthrin, Imidacloprid, Profenofos, Lufenuron, and Fungicides: Cymoxanil, Difenoconazole, Propamocarb). The extractions were made following the QuEChERS protocol (Thermo Scientific, short protocol: AB21891). The extractions were analyzed with ultra-high efficiency liquid chromatography (UHPLC) *Dionex Ultimate 3000* (Thermo Scientific), equipped with a binary gradient bomb (HP G3400RS), an automatic sample (WPS 300TRS), and a thermoset unit for the column (TCC 3000). The LC-MS interface was an electro-nebulization (ESI) and a high-resolution mass spectrometer with an *Orbitrap* detection system of ions. The chromatographic separation was made with the column *Hypersil GOLD Aq* (Thermo Scientific, 100×2.1 mm, 1.9 μm particle size) at 30°C. The mobile phase was A: Aqueous solution 0.2% ammonium formate, and B: acetonitrile with 0.2% ammonium formate. The initial condition of the gradient was 100% A, changing linearly until 100% B (8min). It remained 4 min, with a return to the initial condition in 1 min. The run total time was 13 min, with 3 min of post-run. The mass spectrometer Orbitrap (Exactive Plus, Thermo Scientific) was connected to the electronebulization interface (HESI), operated in positive mode with a capillary voltage of 4.5 kV. Nitrogen was used as drying gas. The mass spectrum was acquired in the mass range 60-900 m/z. The mass detector Orbitrap was calibrated with the certified reference solutions: Ultramark™ 1621 Mass Spec. (AB172435, ABCR GmbH & Co. KG), sodium dodecyl sulfate (L45509, Sigma-Aldrich), and sodium taurocholate hydrated (T4009, Sigma-Aldrich). Compound identification was made using the acquisition mode *Full scan* and the ions extraction corresponding to the pesticides tested [M+H]^+^, mass measuring with accuracy and precision of Δ_ppm_ <0.001, and using a mix standard solution of the pesticides.

For the pesticides detected, we calculated the Hazard Quotient (HQ) as an estimation of the pesticide risk based on the residue concentrations found in the anthers over the LD50 for every molecule (HQ = Concentration of the pesticide residue in the anthers / acute oral LD50 for honey bees, modified from Stoner and Eitzer, 2013). No estimate of pollen consumption was included in this calculation since bees found in the crop are mostly generalists and do not rely on just lulo pollen. Hazard quotients for every pesticide in each farm were summed to estimate a total HQ per farm (Supplementary Table S3). To estimate the contribution of every pesticide to a total hazard in this agroecosystem we calculated an HQ per molecule based on the mean concentration found in the anthers across farms (Table 3).

### 2.4 Bee diversity and flower visitation

To characterize the diversity of the pollinator community in lulo crops, during the dry season, when most of the blooming occurred, we sampled the flower visitors on every farm three times in September, October, and November. Two observers walked for 30 minutes during warm and sunny days from 9:00 to 16:00h, collecting and counting the number of bees observed visiting the flowers. Collected specimens were identified to the lowest level possible depending on the availability of entomological keys and reference material for the country in the bee collection of the Bee Research laboratory (LABUN) at the National University of Colombia in Bogota. We calculated species richness (number of bee species) and abundance (number of bee visits) at every farm.

### 2.5 Pollination experiments and yield

We selected 20 lulo plants in a transect throughout the field to estimate the pollinators’ contribution to fruit production at each farm during the first two weeks of September. Plants were at least 6m apart from each other. For every plant, we chose a pair of closed inflorescences with similar size but in different branches (30cm to 50 cm apart). We bagged one inflorescence using fine organza bags (13×18cm) and left the other open to be visited by the naturally occurring flower visitors. We counted the number of hermaphrodite and staminate flowers to later calculate the fruit set as the number of developed fruits over the number of hermaphrodite flowers. Two weeks later, after all petals had fallen off the flowers, we removed the bags and counted the number of fruits. Three and a half months later, when most of the fruits reached the commercial harvest point, we weighed the fruits and measured the fruit diameter.

### 2.6 Statistical analysis

All the analyses were conducted in R version 3.3.0 (R Core Team, 2019). We examined the effects of landscape composition, pesticide residue, and their interaction on bee diversity and abundance using generalized linear mixed-effects models (GLMM) (Bates et al., 2015) with farm as a random effect to account for nonindependence of samples taken from the same site. However, the variance contributed by farm was very close to zero making the inclusion of the random effect unnecessary (Bolker et al., 2009). We then built models with generalized linear models (GLM) with Poisson error distribution and log link for richness, and quasipoisson and log link for abundance to adjust for overdispersion. To test for spatial autocorrelation of the residuals, we used the Moran’s I test with the function *testSpatialAutocorrelation* from the R package *DHARMa* (Hartig, 2020), and found no autocorrelation for either variable (Richness: *p= 0.085,* Abundance*: p= 0.56).* We also evaluated the effects of richness, abundance, bagged and open treatments, landscape composition, and pesticide residue on fruit set using a GLM with quasibinomial error distribution and logit link function. Quasibinomial error was used to correct for underdispersion. We then obtained the best average model with the *dredge* and *model.avg* functions of the MuMIn package based on Δ2 Quasi Akaike Information Criterion (Barton, 2019). To assess the statistical significance of each explanatory variable and interaction terms of the final models we used Chi-square test for the Poisson fitted model and conditional F tests for the quassipoisson and quasibinomial fitted models with the function *anova.glm* of the stats package (R Core Team, 2019). We used paired t-tests to compare average fruit sets, fruit weights, and fruit diameters among pollination treatments (open and bagged flowers) to determine the contribution of pollinators to lulo yield across farms.

## 3. Results

### 3.1 Lulo crop pollinator community

In this study, we registered 650 bee visits to lulo flowers from 16 bee species. The most common flower visitor was *Tetragonisca angustula* (33.7%, n=216) followed by *Melipona gr. fasciata* (18.9%, n=121), *Paratrigona opaca* (17.7%, n=113) and *Apis mellifera* (11.6%, n=74; Table 1). Bee richness per farm ranged from 1 up to 12 bee species visiting the flowers during the three samplings.

**Table 1.** List of bee species visiting *Solanum quitoense* crops in Casanare, Colombia. The list is organized from the most common visitor presented at the top, to the least common visitor presented at the bottom. ‘n’ represents the total number of visits observed over three samplings dates, each sampling was 30 minutes long with two observers, and ‘%’ the percentage of visits from that one species in relation to the total number of visits observed.

| Tribe | Species | n | % |
| --- | --- | --- | --- |
| Meliponini | *Tetragonisca angustula* | 216 | 33.75 |
| Meliponini | *Melipona gr. fasciata* | 121 | 18.91 |
| Meliponini | *Paratrigona opaca* | 113 | 17.66 |
| Apini | *Apis mellifera* | 74 | 11.56 |
| Bombini | *Bombus transversalis* | 31 | 4.84 |
| Euglossini | *Euglossa sp. (Females)* | 21 | 3.28 |
| Euglossini | *Eulaema cingulata* | 14 | 2.19 |
| Euglossini | *Eulaema mocsaryi* | 14 | 2.19 |
| Augochlorini | *Augochloropsis sp. 1* | 9 | 1.41 |
| Augochlorini | *Augochlora sp.1* | 8 | 1.25 |
| Bombini | *Bombus excellens* | 8 | 1.25 |
| Meliponini | *Trigona fulviventris var guianae* | 7 | 1.09 |
| Meliponini | *Plebeia sp.1* | 1 | 0.16 |
| Meliponini | *Scaptotrigona sp.1* | 1 | 0.16 |
| Eucerini | *Thygater aethiops* | 1 | 0.16 |
| Eucerini | *Thygater analis* | 1 | 0.16 |

### 3.2 Pesticide results

For the pesticide detections, we found 5 out of the 9 molecules tested in the anthers. The two most frequent pesticides found were the fungicide Propamocarb in 100% of the farms, ranging from 1.3 to 399.5 μg/kg, and the insecticide imidacloprid in 80% of the farms ranging from 0.6 to 13063 μg/kg. The contribution of every pesticide to the total HQ shows that imidacloprid contributes 99.34% to the total hazard being the main molecule generating a risk to pollinators in this cropping system (Table 2).

**Table 2.**
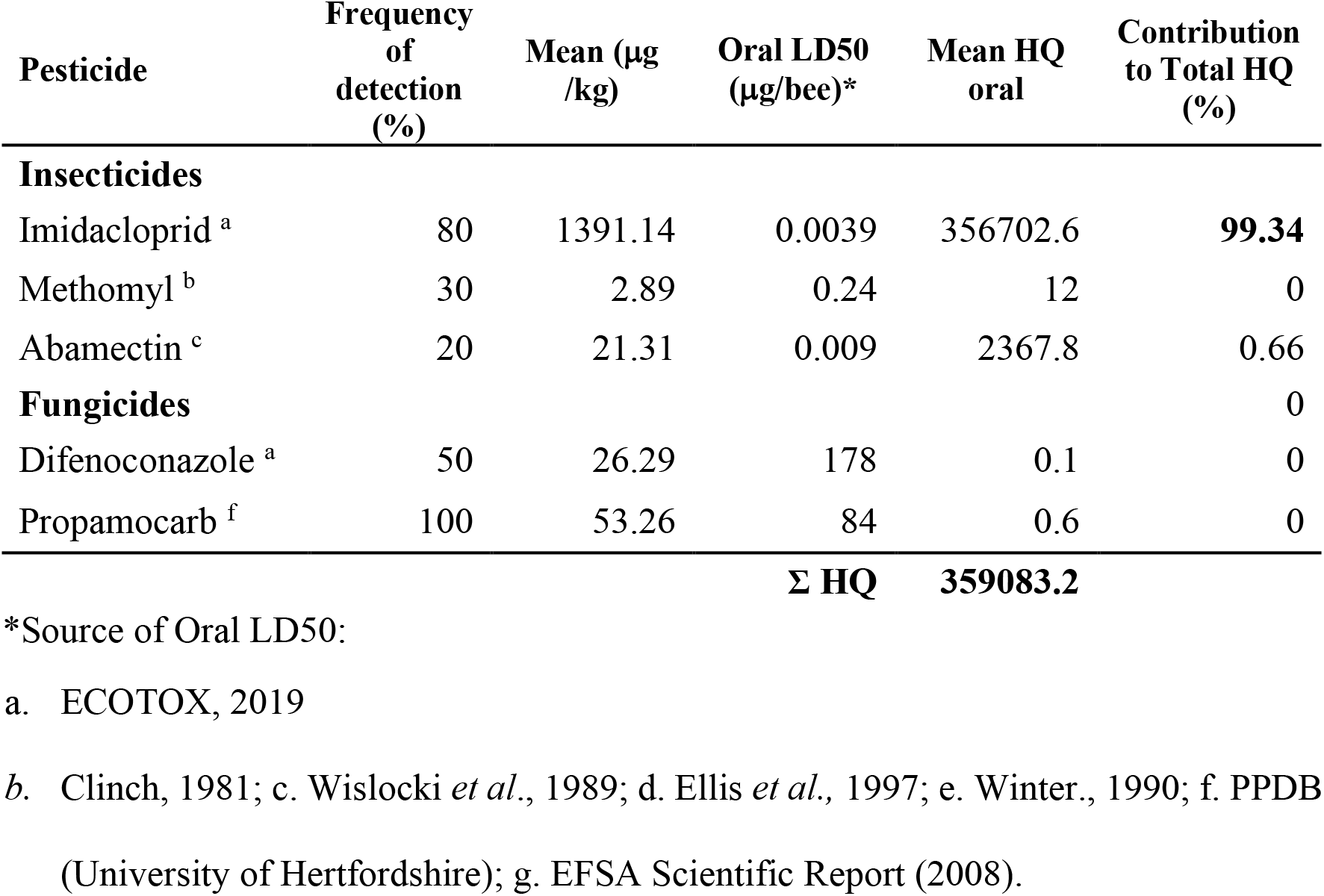
Hazard quotients (HQ) calculated for the pesticides detected in anthers of *S. quitoense* crops in Casanare, Colombia, and the contribution to a Total hazard quotient for the cropping system in the region.

### 3.3 Landscape and pesticide effects on bee richness

Overall, as the proportion of natural habitat increased bee richness increased (*X^2^_(1,28)_= 6.9, p-value=0.0086, Fig.1a*), and as the HQ per farm increased bee richness decreased (*X^2^_(1,27)_=11.06,p-value* <0.001*, Fig.1b*). Further, these two factors were interacting (*X^2^_(1,26)_ = 5.54, p-value= 0.0186, Fig.2a*). The negative effect of an increased HQ on bee richness was less detrimental in farms with higher proportions of natural habitat. For example, for farms No. 1 and No. 5 with similar HQ levels (828,5 and 743,7 respectively; Supplementary Table S3) the richness in farm No.5 (10 species) was twice as large as in farm No.1 (5 species), and this difference can be explained by the higher proportion in natural habitat in farm No.5 (Farm 1: 29,4%, Farm 5: 62,3%). But this buffer effect is lost in farms where the HQs are very high (>10,000; *Fig. 2a*).

**Figure 1.**
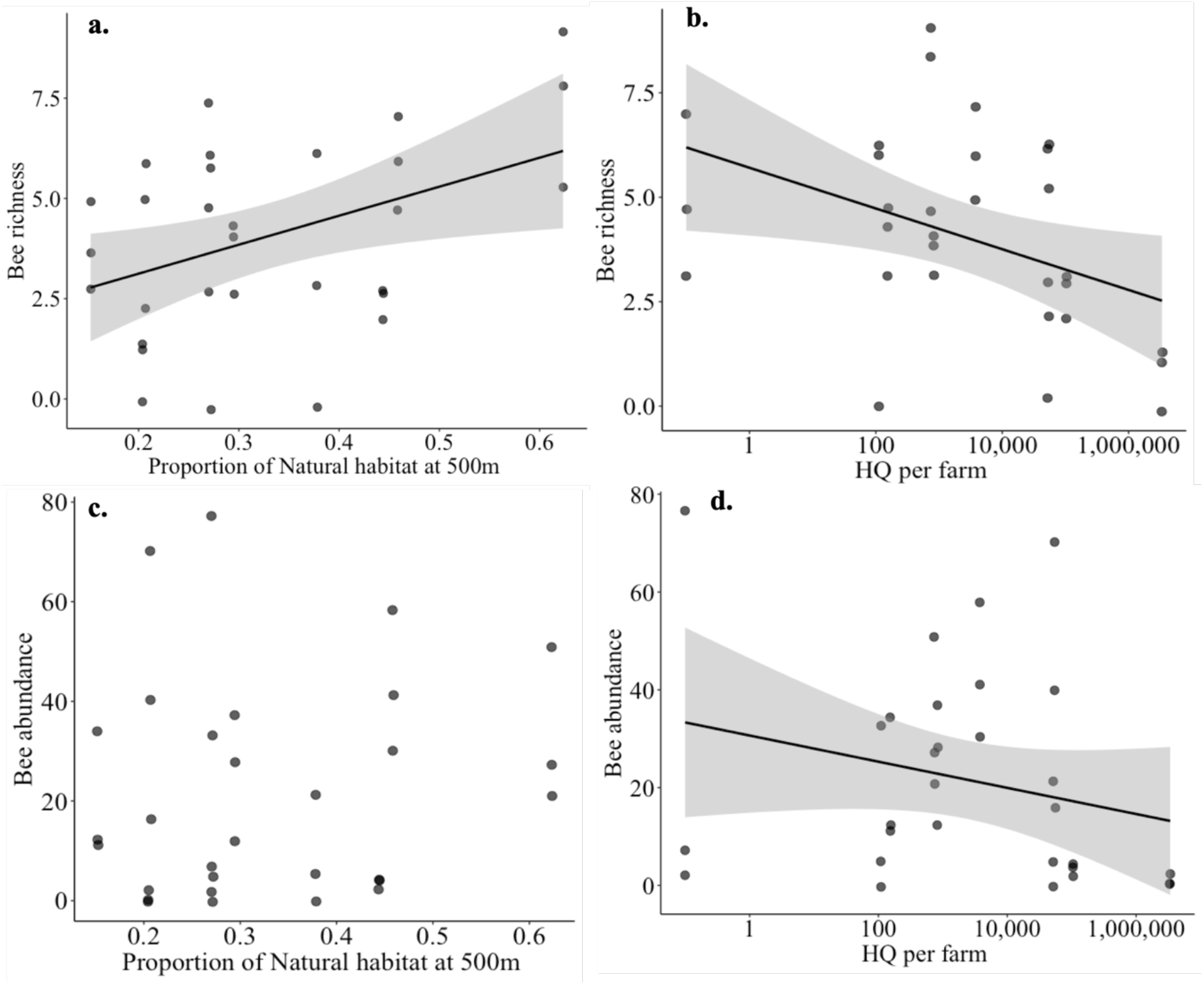
**(a-b)** Bee richness and **(c-d)** abundance in relation to the proportion of natural habitat at 500m around the *S. quiotense* crops, and the hazard quotient (HQ) calculated for the pesticide residues found in the anthers (x axis for HQ is log-transformed). Lines represent a significant regression between the two factors (*p<0.005*) and the grey areas are the standard errors.

**Figure 2.**
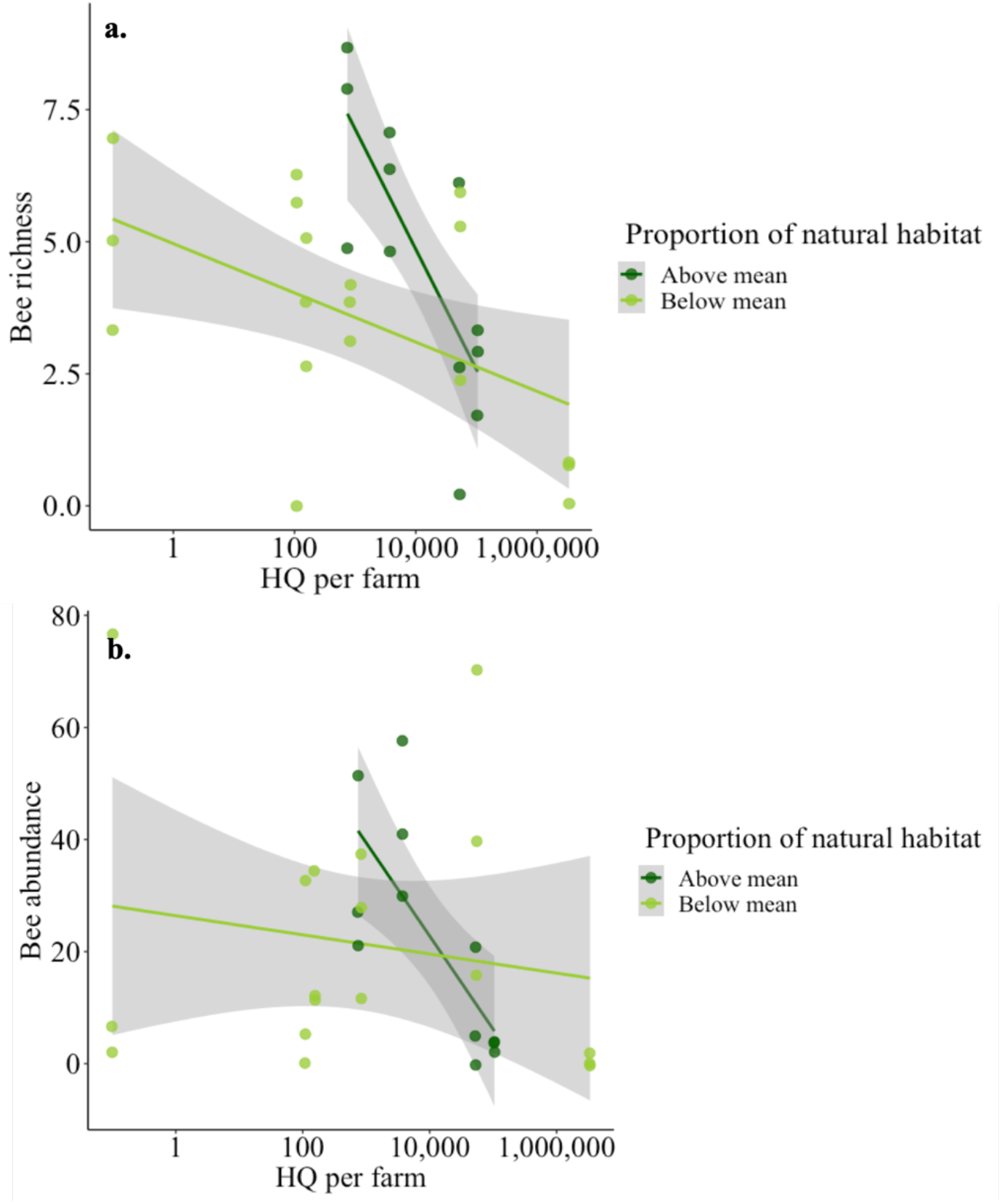
**(a)** Bee richness and **(b)** bee abundance in relation to the Hazard Quotient (HQ) calculated per farm (x axis log-transformed) for two different levels of the proportion of natural habitat at 500m in *S. quiotense* crops (Dark green: above mean, light green: below mean, mean=0.33). Natural habitat categories were only used for graphical purposes to depict the interaction effect, but they were not used in the regression model analysis.

### 3.4 Landscape and pesticide effects on bee abundance

We found no significant effect of natural habitat proportion on bee abundance (*F_(1,28)_=0.7625, p-value=0.390, Fig.1c*), while as HQ per farm increased bee abundance decreased (*F_(1,27)_=8.4243, p-value=0.007), Fig.1d*). However, there was a significant interaction effect (*F_(1,26)_=8.80, p-value=0.0063), Fig.2b*), where a higher proportion of natural habitat attenuates the negative effects of pesticides on bee abundance, but once again this effect is lost in farms where HQs are above 10,000 (*Fig. 2b*).

### 3.5 Yield

We found a mean number of 11 (+/-2.2) flowers per inflorescence, 71% staminate, and 29% hermaphrodite flowers. For all the fruit yield variables, the open pollination treatment had a higher yield than the bagged treatment. The mean fruit set was 4% for the bagged treatment (bagged inflorescences), and 61% for open-pollinated treatment (*t = −7.982, df = 9.5087, p-value<0.001, Fig.3a*). The mean fruit weight was 61.8 g for bagged flowers and 86.2 g for open pollination (*t = 13.814, df = 23.117, p-value<0.001, Fig. 3b*). Fruit diameter also differed among treatments: the mean for the bagged treatment was 4.37cm and 5.49 cm for open pollination (*t = 8.8456, df = 21.324, p-value<0.001, Fig. 3c*).

**Figure 3.**
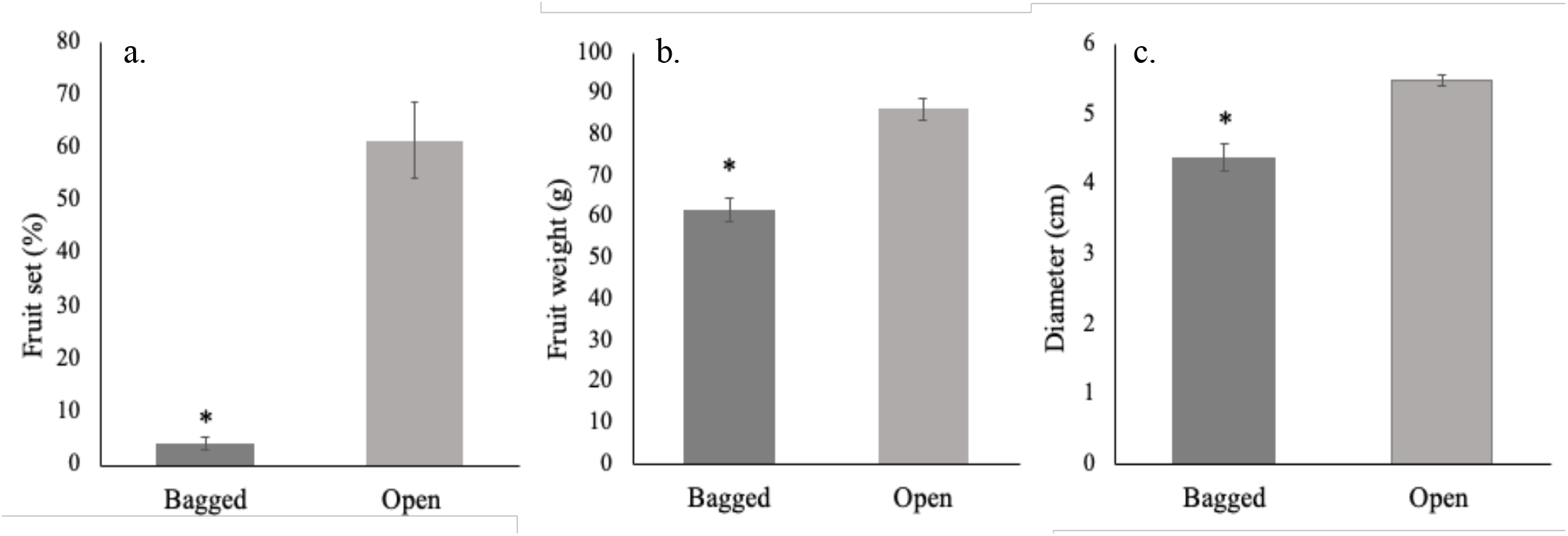
Average fruit set **(a),** fruit weight **(b),** and fruit diameter **(c)** (+/− 1 SE) of bagged and open pollinated *S. quitoense* fruits across farms. (*) denotes significant differences among treatments.

The best supported model for fruit set (fruit set ~ open/bagged treatment + richness + HQ+ Natural habitat) showed that bee richness (*F_(1,367)_=4.97, p-value=0.0262, Fig.4*) and open/bagged treatments had significant effects on fruit set (*F_(1,370)_= 292.5, p-value=<0.001*). While HQ had a weak effect (*F_(1,368)_= 2.928, p-value=0.087*) and the proportion of natural habitat was not significant (*F_(1,369)_ = 0.0087, p-value=0.925*).

**Figure 4.**
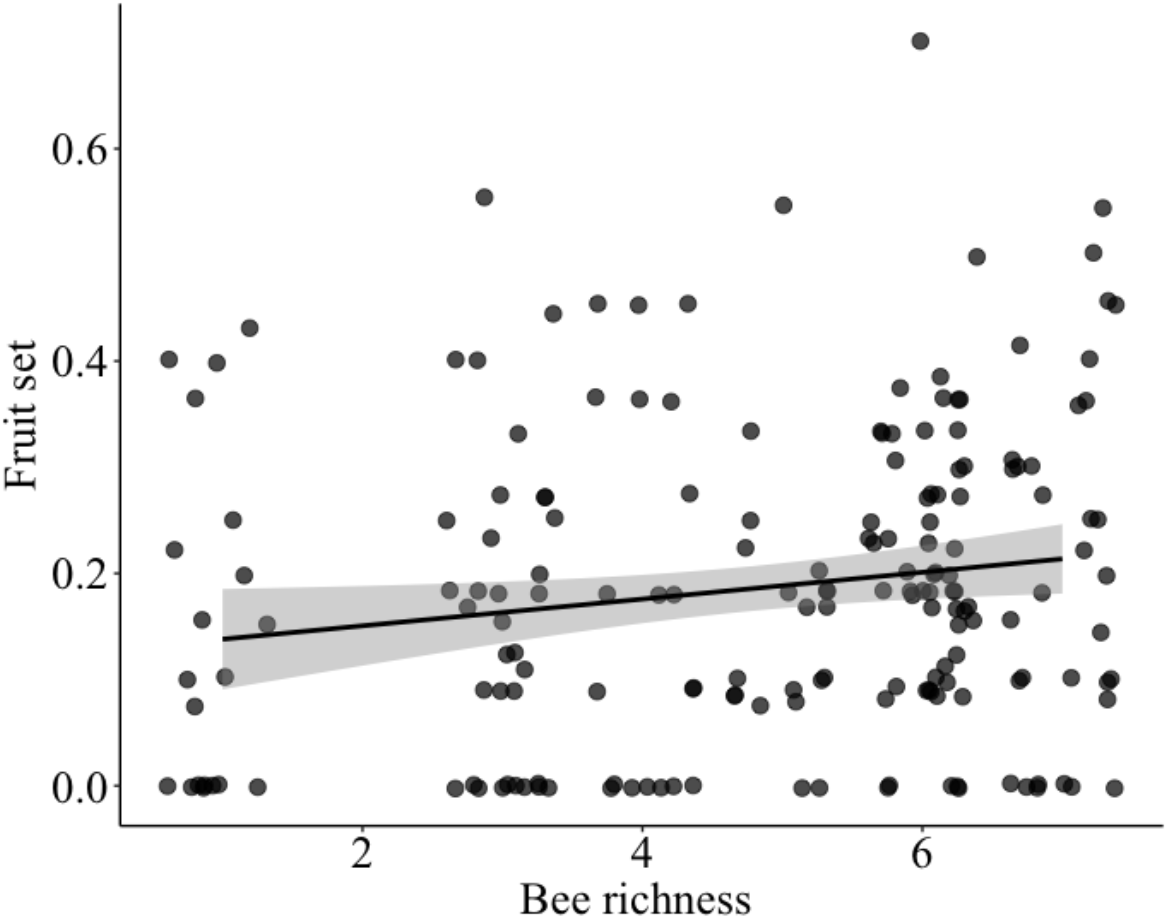
Fruit set in relation to bee richness in *S. quiotense* crops. Line represents a significant regression between the two factors (*p<0.005*) and the grey area the standard error.

## 4. Discussion

We assessed the landscape composition and pesticide residue effects on bee diversity, abundance, and pollination services in *S. quitoense*. Our results provide evidence that the natural habitat loss due to the expansion of pastures for cattle ranching reduces bee diversity in this tropical cropping system. This may occur because the pollinator community requires floral and nesting resources that are more abundant in natural habitats than grasslands. In tropical forests, it has been shown that bee community composition changes with forest fragmentation and traits like tree-cavity nesting in Meliponini bees or loss of particular plant groups (*i.e.* epiphytes for Euglossini) are determinant in those changes (Allen et al., 2019; Brosi et al., 2007). We found a significant decline in richness but not in abundance, suggesting that in farms with less natural habitat in the landscape, vulnerable species are being lost but common species could be taking advantage of these resources provided by the crop.

From the five pesticides detected, the fungicide propamocarb and the insecticide imidacloprid were the most prevalent. It should be noted that both products are systemic, which means that although they are applied to the soil or foliage in this crop, they can reach the floral organs, as shown by our results. Propamocarb is considered low to moderately toxic to honey bees (EFSA, 2006), and in this study, we did not find any impact on the bee community. However, studies on sub-lethal effects and long-term effects are required. In contrast, imidacloprid is a highly toxic molecule for bees (EFSA, 2012), and the concentrations found were extremely high, far above any other concentration previously reported in flowers for any crop (Blacquière et al., 2012). In the region, farmers attribute the excessive use of insecticides to the control of a fruit borer moth (*Neoleucinodes elegantalis*), which oviposits on newly formed fruits. Complementary studies must consider the level of this pest in different farms which can explain the high quantity of pesticides found. Additionally, it has been described that smallholder growers in Colombia tend to over- or mis-use use pesticides in solanaceous crops (Bojacá et al., 2013; Feola and Binder, 2010).

In this field study, we show that bee richness and bee diversity decreased as the anther HQ per farm increased. Additionally, based on the toxicity and the frequency of the pesticides found in the samples, imidacloprid is the pesticide that represents the greatest risk for bees, as has been shown in studies for other regions of the world (Sanchez-Bayo and Goka, 2014). The same negative effect has been demonstrated in apple orchards in Wisconsin, where abundance and richness of *Lasioglossum* species decreased with the increased use of imidacloprid (Mallinger et al., 2015). Our approach of testing the actual residuality in the anthers provides a field-realistic estimation of the pesticide exposure generated by this crop, given that *S. quitoense* does not produce any nectar. However, while the HQ based on LD50s for honey bees could predict effects on the whole pollinator community, specific effects must be studied since it has been demonstrated that different bee species vary in their susceptibility to pesticides. Bumble bees and stingless bees in particular, have shown greater susceptibility to certain pesticides compared to honey bees (Cresswell et al., 2014; Del Sarto et al., 2005).

Our results provide new evidence about the interactive effects between landscape composition and pesticide exposure, showing that natural habitat area surrounding crops can help to mitigate negative pesticide effects on bee communities. However, this buffer effect is present until certain levels of HQ, showing that very high concentrations of pesticides are deleterious in any landscape. This might be explained by the fact that most of the bees visiting this crop are generalists, and depending on their diet, the pesticide toxicity can be diluted. A model created for honey bees foraging in landscapes with oilseed rape showed that a significant pesticide dilution can occur when alternative resource patches are equal or more attractive than the focal crop and these are close and abundant (Baveco et al., 2016).

We found 16 bee species from 6 tribes visiting lulo flowers. *Tetragonisca angustula* and *Paratrigona opaca* were two of the most frequent visitors, and although they do not perform sonication, they were observed removing pollen through the anther pores using their glossa and hind legs. How efficient they are at transporting pollen to the stigma is still in question.

On the other hand, we observed 10 species sonicating the flowers, among them *M. gr. fasciata* was the most abundant. Other species of the genus *Melipona* have been previously reported as efficient pollinators of Solanaceous crops such as tomato and eggplant (Bartelli and Nogueira-Ferreira, 2014; Nunes-Silva et al., 2013). A study assessing the effects of imidacloprid consumption in bumble bees pollinating tomatoes showed that imidacloprid affected the probability of performing sonication behavior in flowers (Switzer and Combes, 2016). This might suggest that pesticides in lulo are not just affecting bee diversity but also their capacity to release the pollen during visitation.

We found a 51% reduction in fruit set when flower visitors were excluded, showing a high pollinator dependency according to the Klein *et al.* (2007) classification (40 to 90% reduction: high dependency). We also found a 39% decrease in fruit weight and 25% in fruit diameter without bees demonstrating their contributions to crop yield. These results are similar to those found on artificial cross-pollination treatments in greenhouses for lulo (Almanza, 2007; Messinger et al., 2016). We did not find significant effects of landscape composition, or pesticides on fruit set while bee richness was significant. This suggest that these factors do not affect the yield directly, but through the decrease of the pollinators.

The results of this study can help farmers, land planners, and local institutions to make decisions towards pollinators protection measures before the reduction of bees is exacerbated, leading to more pronounced effects on crop productivity. This study supports the protection of natural areas surrounding crops as a strategy to promote crop pollinator communities. Based on the wide variation and the high amounts of imidacloprid found, it is also necessary to understand the factors that are driving these applications in crops and develop strategies to reduce them. In addition, restoration and the implementation of local diversification strategies might be established to aid bees to overcome the risks associated with pesticides.

## Acknowledgments

A special thank you to the participating growers who provided the field sites for this study. Thanks to David Sossa for invaluable field assistance. Also thank you to Rodulfo Ospina, Diego Guevara, Jorge Diaz, and Joanna Jaramillo for bee identifications. We are grateful to Scott McArt, Bryan Danforth, Beatriz Ramirez, and Fiona Rodgerson for helpful revisions to earlier versions of the manuscript. D.O. was funded by Fulbright - COLCIENCIAS (Colombian Administrative Department of Science, Technology, and Innovation) through a doctorate scholarship. Fieldwork for this study was funded by the Richard Bradfield Research Award and the Grace H. Griswold Fund, both from the College of Agriculture and Life Sciences at Cornell University.

## Conflict of interest

The authors declare no conflict of interest.

## Supplementary information

**Figure S1.**
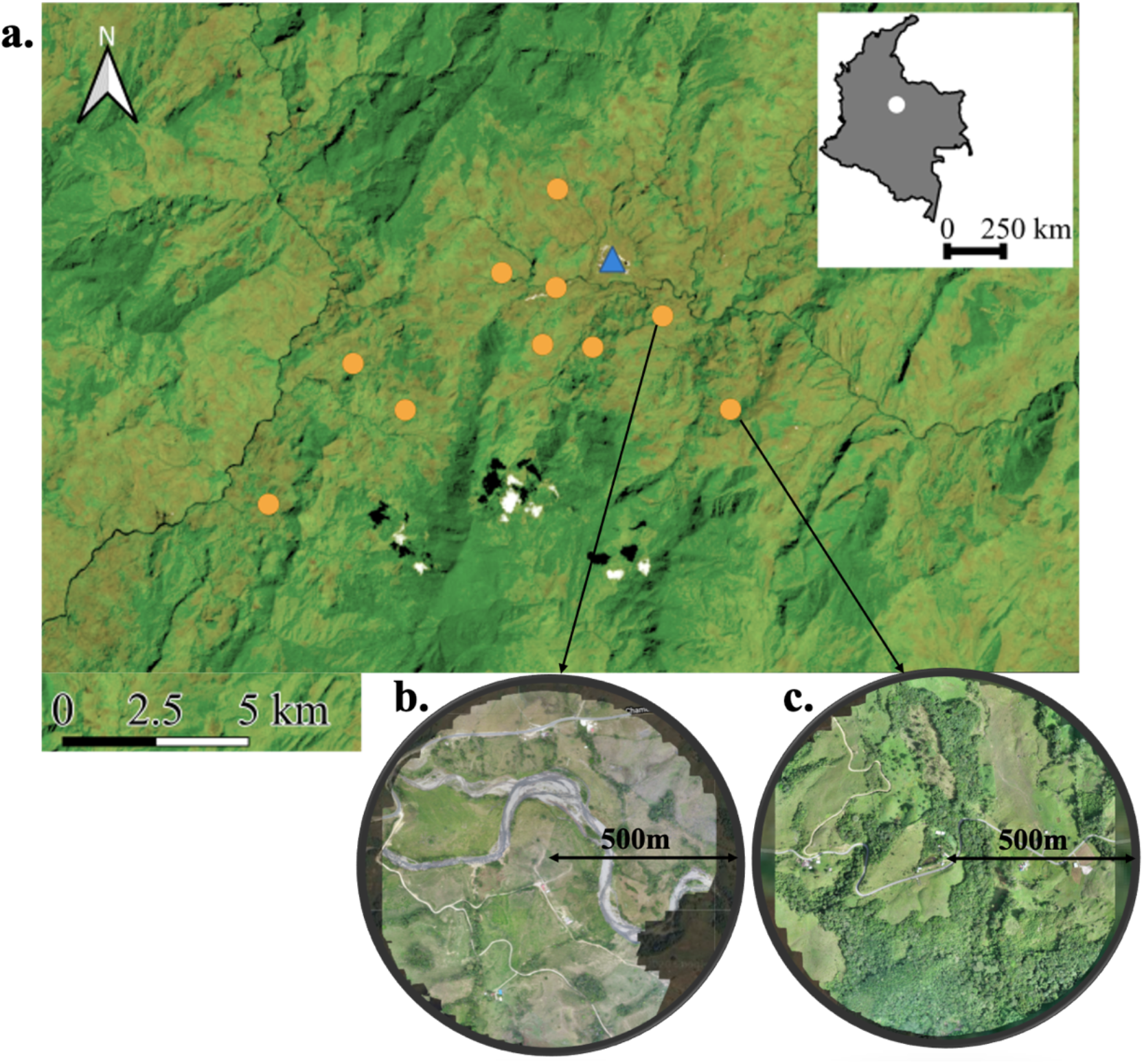
a. Satellite image of the study area, orange dots are the locations of the sampled farms and the blue triangle is the urban area of Chameza. b. Orthophoto constructed with drone pictures from a farm within a simple landscape at 500m (72% Pastures, 15% Natural habitat, 13% Lulo crops) and c. Orthophoto constructed with drone pictures from a farm within a complex landscape at 500m (b. 37% Pastures, 62% Natural habitat, 0.3% Lulo crops).

**Table S2.**
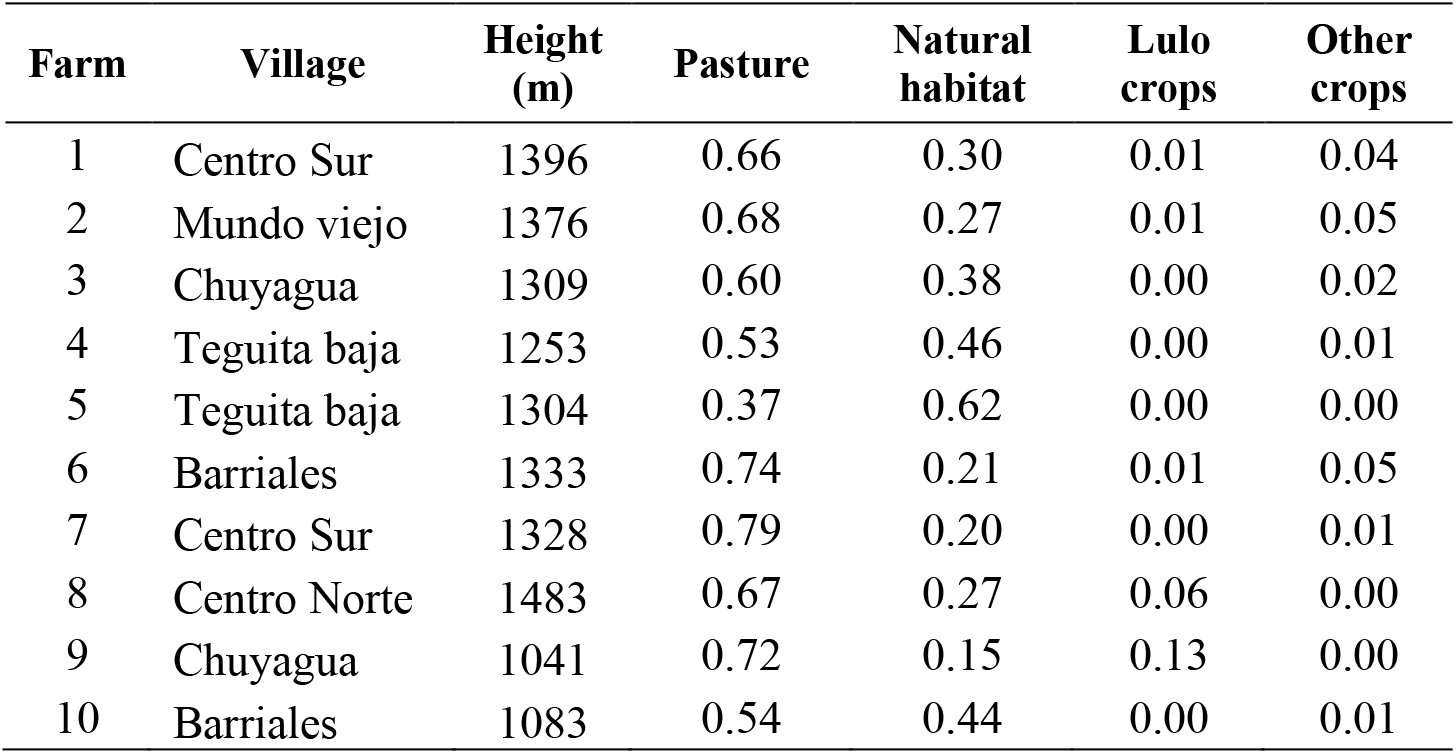
Location, height, and landcover type proportions of every farm in the study at 500m around the lulo crop fields.

**Table S3.**
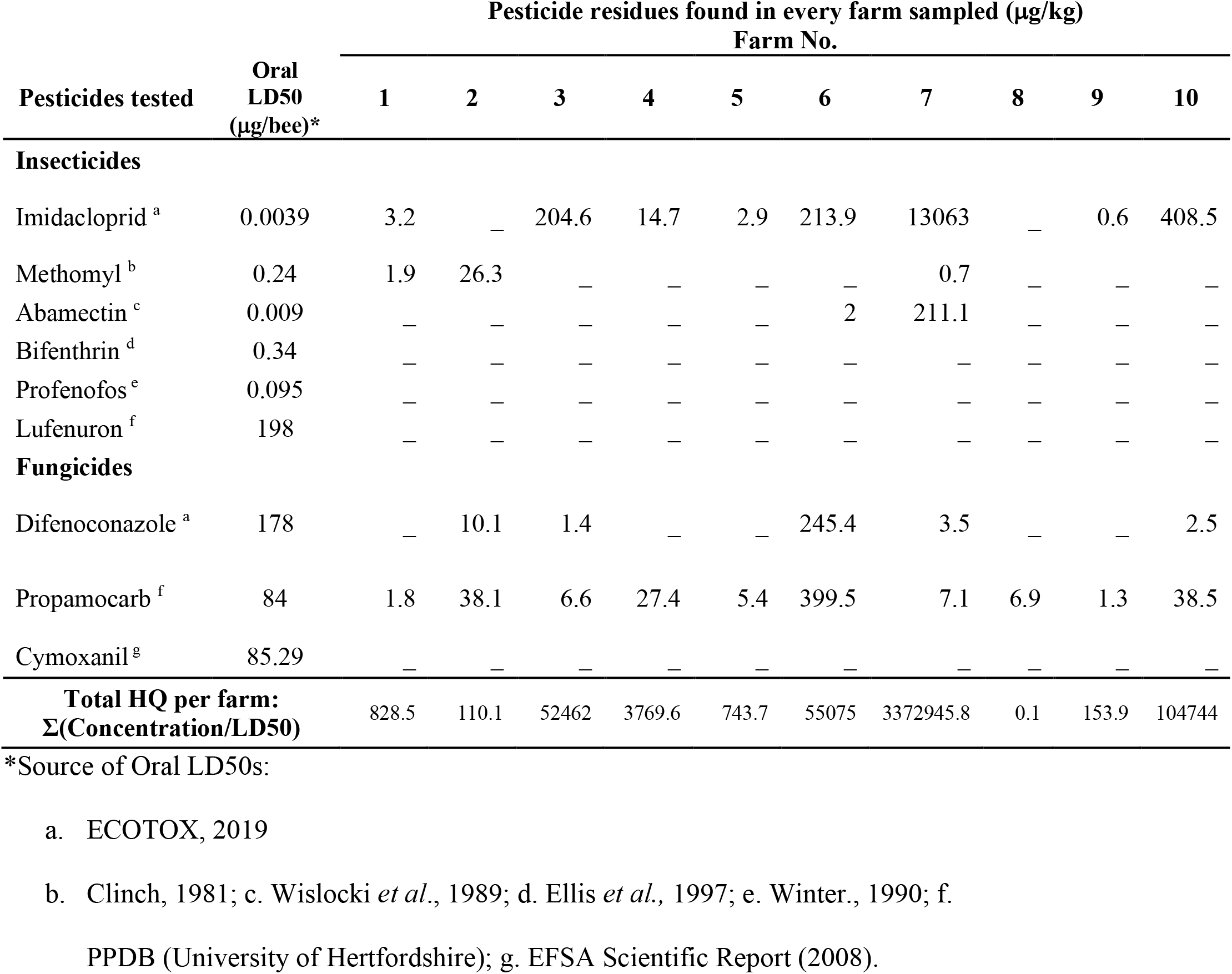
Concentrations of nine pesticides tested in anthers of *S. quitoense* crops in each of the 10 farms included in the study in Casanare, Colombia. The minimum level of quantification was 0.1 μg/kg, while the minimum level of detection was 0.05μg/kg. (_) represents when the compound at the farm was not detected. The last row shows the total hazard quotients (HQ) calculated per farm based on honey bee oral LD50 values from the literature.

| Pesticides tested | Oral<br>LD50<br>(µg/bee)* | Pesticide residues found in every farm sampled (µg/kg) |  |  |  |  |  |  |  |  |  |
| --- | --- | --- | --- | --- | --- | --- | --- | --- | --- | --- | --- |
|  |  | Farm No. |  |  |  |  |  |  |  |  |  |
|  |  | 1 | 2 | 3 | 4 | 5 | 6 | 7 | 8 | 9 | 10 |
| Insecticides |  |  |  |  |  |  |  |  |  |  |  |
| Imidacloprid <sup>a</sup> | 0.0039 | 3.2 | — | 204.6 | 14.7 | 2.9 | 213.9 | 13063 | — | 0.6 | 408.5 |
| Methomyl <sup>b</sup> | 0.24 | 1.9 | 26.3 | — | — | — | — | 0.7 | — | — | — |
| Abamectin <sup>c</sup> | 0.009 | — | — | — | — | — | 2 | 211.1 | — | — | — |
| Bifenthrin <sup>d</sup> | 0.34 | — | — | — | — | — | — | — | — | — | — |
| Profenofos <sup>e</sup> | 0.095 | — | — | — | — | — | — | — | — | — | — |
| Lufenuron <sup>f</sup> | 198 | — | — | — | — | — | — | — | — | — | — |
| Fungicides |  |  |  |  |  |  |  |  |  |  |  |
| Difenoconazole <sup>a</sup> | 178 | — | 10.1 | 1.4 | — | — | 245.4 | 3.5 | — | — | 2.5 |
| Propamocarb <sup>f</sup> | 84 | 1.8 | 38.1 | 6.6 | 27.4 | 5.4 | 399.5 | 7.1 | 6.9 | 1.3 | 38.5 |
| Cymoxanil <sup>g</sup> | 85.29 | — | — | — | — | — | — | — | — | — | — |
| Total HQ per farm:<br>Σ(Concentration/LD50) |  | 828.5 | 110.1 | 52462 | 3769.6 | 743.7 | 55075 | 3372945.8 | 0.1 | 153.9 | 104744 |

\*Source of Oral LD50s:
a. ECOTOX, 2019
b. Clinch, 1981; c. Wislocki *et al.*, 1989; d. Ellis *et al.*, 1997; e. Winter., 1990; f.
PPDB (University of Hertfordshire); g. EFSA Scientific Report (2008).

## Notes

### Competing Interest Statement

The authors have declared no competing interest.

